# driveR: A Novel Method for Prioritizing Cancer Driver Genes Using Somatic Genomics Data

**DOI:** 10.1101/2020.11.10.376707

**Authors:** Ege Ülgen, O. Uğur Sezerman

## Abstract

Cancer develops due to “driver” alterations. Numerous approaches exist for predicting cancer drivers from cohort-scale genomic data. However, methods for personalized analysis of driver genes are underdeveloped.

In this study, we developed a novel personalized/batch analysis approach for driver gene prioritization utilizing somatic genomic data, called driveR. Combining genomic information and prior biological knowledge, driveR accurately prioritizes cancer driver genes via a multi-task learning model.

Testing on 28 different datasets, this study demonstrates that driveR performs adequately, outperforms existing approaches, and is an accurate and easy-to-utilize approach for prioritizing driver genes in cancer genomes. driveR is available on CRAN: https://cran.r-project.org/package=driveR.

## 1. Background

Cancer develops due to changes that have occurred in the DNA sequence of the genomes of cancer cells, somatic mutations acquired during the lifetime of an individual [1]. Cancer genomes contain large numbers of somatic mutations, but most are “passengers” that emerge simply as a result of genome instability during cancer progression and do not contribute to cancer development, and a small proportion are “drivers” that are implicated in oncogenesis [2-4]. Identification of molecular cancer driver genes is critical for personalized oncology as accurate identification of personalized driver genes will result in precise diagnosis and will allow the clinicians to possibly define personalized therapeutic targets [5, 6]. Several computational batch analysis approaches, extensively reviewed by Tokheim et al. and Cheng et al. [7, 8], have been developed to identify cancer driver genes. Some notable approaches for batch analysis include MuSiC [9], MutSigCV [10], MutPanning [11], MEMo [12], Hierarchical HotNet [13], TieDie [14], DriverNet [15], CaDrA [16] and OncodriveFML [17]. MuSiC uses the significance of a higher than expected rate of mutations, pathway mutation rate, and correlation with clinical features to detect drivers. MutSigCV investigates the mutational significance of genes by identifying genes that were mutated more often than expected by chance given background mutation processes. MutPanning uses deviation of mutational context from characteristic contexts around passenger mutations in addition to traditionally-used features (e.g. a higher than expected mutation rate) for driver gene identification. MEMo tries to detect small subnetworks of genes in the same pathway, exhibiting internal mutual exclusivity patterns. Hierarchical HotNet incorporates knowledge from protein-protein interaction networks (PINs) to find a hierarchy of altered subnetworks, containing frequently mutated genes. TieDie incorporates PIN and mRNA expression data to find overlapping subnetworks exhibiting a high degree of mutation and expression values using heat diffusion. DriverNet tries to detect driver genes via their effect on mRNA expression networks, by identifying a set of genes with mutations/copy-number alterations that are linked to genes with deregulated expression in a given PIN. CaDrA uses a step-wise heuristic search approach to identify functionally relevant subsets of genomic features, maximally associated with a specific outcome of interest. OncodriveFML aims to detect driver genes by analyzing the functional impact bias of observed somatic mutations.

Personalized driver gene prioritization is essential for numerous reasons: (i) it is important to identify true driver genes for the patient as some patients have alterations in many known driver genes, (ii) there may be a need to identify putative driver genes in patients without any alteration in any known driver gene, (iii) as the number of therapies that can be administered at the same time is limited due to toxicity and adverse events [18, 19], there is a need to prioritize driver genes for the patient. Although there are several approaches for identifying cancer driver genes in cohort-scale genomics data, personalized driver gene identification approaches are still underdeveloped. Some methods that operate on the data of a single patient to identify and rank patient-specific driver genes have been developed: DawnRank uses a PageRank algorithm to rank mutated genes according to the effect on expression deregulation of downstream differentially-expressed genes (DEGs) in a directed PIN [20]. The single-sample controller strategy (SCS) aims to identify a set of mutated genes that are linked to downstream DEGs in a directed PIN [21]. iCAGES utilizes a statistical framework to identify driver variants by integrating contributions from coding, non-coding, and structural variants and identifies driver genes by combining genomic information and prior biological knowledge [22]. PRODIGY analyzes the expression and mutation profiles of the patient along with data on known pathways and PIN to quantify the impact of each mutated gene on every deregulated pathway using the prize-collecting Steiner tree model [23].

As stated above, most approaches for driver gene prioritization are batch-analysis-based. Furthermore, most approaches utilize only genomics and/or transcriptomics data without exploiting prior biological knowledge. To improve on existing approaches, we aimed to develop a novel driver gene prioritization approach that utilizes somatic genomics information incorporating prior biological knowledge, which we called driveR. We developed driveR intending to establish an accurate and reliable method for driver gene prioritization. The approach allows for personalized or batch analysis of genomics data for driver gene prioritization by combining genomic information and prior biological knowledge. As features, driveR uses coding impact metaprediction scores, non-coding impact scores, somatic copy number alteration scores, hotspot gene/double-hit gene condition, Phenolyzer [24] gene scores, and memberships to cancer-related Kyoto Encyclopedia of Genes and Genomes (KEGG) [25] pathways. It uses these features to estimate cancer-type-specific probabilities of being a driver for each gene using the related task of a multi-task learning model. In this article, we demonstrate that our approach can help increase the accuracy of cancer driver gene detection and prioritization, and therefore, facilitate precise diagnosis, personalized therapy, and overall result in better clinical decision-making.

## 2. Methods

### 2.1. Coding variant impact metaprediction

Initially, we fitted coding impact metapredictor models to assign an estimated probability of damaging impact for each somatic variant. This model was later used to generate a feature for each gene in the multi-task learning classification model. We ensured that the genomic coordinates used in this study are hg19.

For training the coding impact metapredictor models, a benchmarking dataset from the Martelotto et al. study [26] was obtained. All mutations within the dataset were annotated using ANNOVAR [27] with the 12 impact predictors using dbNSFP v3.0 [28]: SIFT [29], PolyPhen-2 [30] (HumDiv scores), LRT [31], MutationTaster [32], Mutation Assessor [33], FATHMM [34], GERP++ [35], PhyloP [36], CADD [37], VEST [38], SiPhy [39] and DANN [40]. Variants with any missing predictor scores were excluded. Additionally, variants with the label “uncertain” were excluded, hence, the final dataset consisted of 135 “neutral” and 814 “non-neutral” variants.

Before training the models, pairwise Pearson correlations of all variables (including the outcome variable) were investigated to determine any collinearity issue and to evaluate the predictive strength of the predictors (Figure S1A). The correlogram revealed that each of the individual scores provides a different facet of information for a coding variant. Additionally, violin plots of all 12 predictors by outcome status (i.e., “neutral” or “non-neutral”) were visualized for investigating the distributions of the two status labels and detect any outliers (Figure S1B). The violin plots revealed that the distributions of scores in “neutral” and “non-neutral” variants were different. There were no noticeable outliers.

The final dataset was split into training and test datasets by randomly selecting 75% of each “neutral” and “non-neutral” variants as the training set and the remaining as the test set.

In total, 6 different classification models were trained and evaluated: logistic regression, naïve Bayes, support vector machine (SVM) with linear kernel, SVM with radial kernel, random forest, and gradient boosting machine. Each of the 6 classification models was trained using the training dataset using 10-fold 3-times-repeated cross-validation, maximizing the Area Under Curve (AUC) metric, and tested on the test dataset. Figure S2 shows the receiver operating characteristic (ROC) curves and AUC values for each of the 6 classification models in the training, test, and validation datasets.

The model with the best performance in all of the datasets was the random forest classification model and this model was used as the coding variant impact metapredictor.

### 2.2. Driver gene prioritization

The overview diagram of our approach to prioritize driver genes is presented in Figure 1. We again ensured that the genomic coordinates used in this study are hg19. For prioritizing cancer driver genes in a cancer-type-specific context, a multi-task learning (MTL) classification model was trained. For this purpose, initially, all somatic mutation and somatic copy number alteration (SCNA) data from The Cancer Genome Atlas (TCGA) program available on the International Cancer Genome Consortium (ICGC) data portal [41] was obtained. This data consisted of genomics data for 21 different cancer types (Supplementary Table 1). Genomics data from each of these different cancer types were randomly split into training (75% of patients) and test (25% of patients) data. Additional datasets for testing were obtained for 4 different cancer types from cBio Cancer Genomics Portal [42] (namely, BRCA-

**Figure 1.**
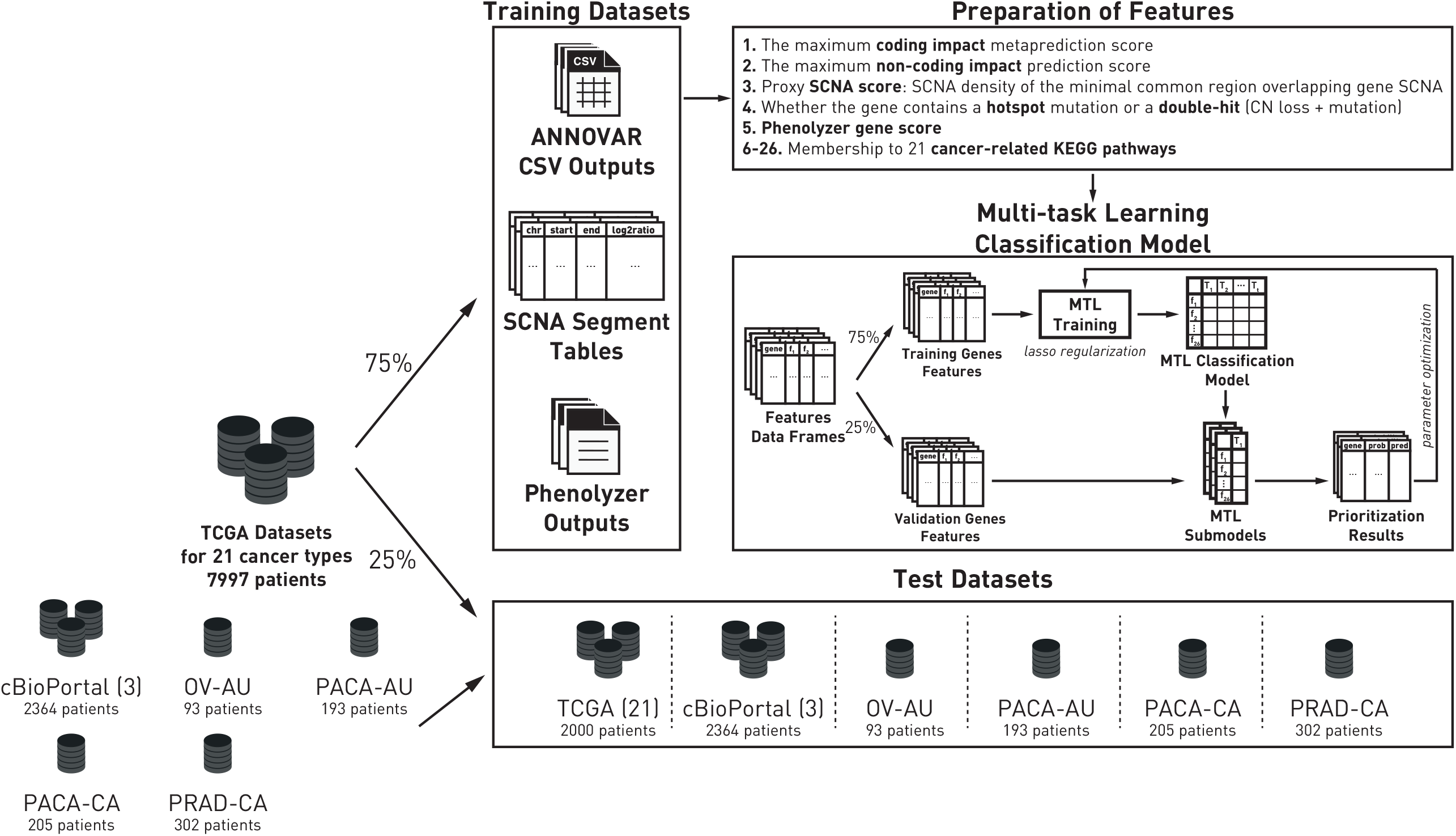
The overview of the driveR approach for driver gene prioritization.

METABRIC [43], COAD-CPTAC [44] and LUAD-ONCOSG [45]) and 4 different cancer types from ICGC (namely OV-AU, PACA-AU, PACA-CA, and PRAD-CA). “True positive” driver genes were defined as the 723 experimentally-validated driver genes curated by the Cancer Gene Census (CGC, v92) [46]. Using ANNOVAR annotations, SCNA tables, and Phenolyzer gene scores, 26 features were generated:

### The maximum coding variant impact score

The maximal coding variant metaprediction score for each gene was used as a feature in the driver gene classification model.

### The maximum non-coding variant impact score

For non-coding variants, the Phred-scaled CADD scores were used. The maximal score for each gene was used as a feature in the driver gene classification model.

### Proxy SCNA score

To score gene-level SCNA events, firstly, the gene-level SCNA events were determined using segment-level SCNA data. If a segment overlapped at least 25% (the default threshold value) of the gene, the gene-level SCNA event was called. The log_2_ ratio for any gene was determined as the ratio with the maximum |log_2_ ratio| value among all segments overlapping the gene. Genes on sex chromosomes were excluded. Gene-level SCNA events with |log_2_ ratio| < 0.25 (the default threshold value) were also excluded.

A Minimal Common Region (MCR) is defined as the minimal region of copy number amplifications or deletions representing a common genomic alteration across the examined cancers [47]. We obtained pan-cancer MCR data from Kim et al. [48] who analyzed chromosomal aberrations in 8000 cancer genomes. Genes overlapping an MCR region with the same direction of SCNA event (amplification or deletion) were assigned the SCNA density (SCNA / Mb) of the MCR. This SCNA density was used as a feature in the driver gene classification model.

### Hotspot or double-hit gene condition

Genes containing hotspot mutations (used as an indication of oncogenes) were determined using the Catalogue of Somatic Mutations in Cancer (COSMIC) [49] v92 occurrence annotations. A mutation with an occurrence greater than 5 (the default threshold) was defined as a hotspot mutation. Additionally, genes with a non-synonymous mutation and a homozygous copy number loss (defined as |log_2_ ratio| < -1) was defined as a double-hit gene (used as an indication of tumor suppressor genes). Whether or not a given gene was a hotspot gene or a double-hit gene was used as a feature in the driver gene classification model.

### Phenotype based gene analyzer score

Phenotype Based Gene Analyzer (Phenolyzer) [24] is a tool for phenotype-based prioritization of candidate genes using prior biological knowledge and phenotype information. All genes from genomics data were used as an input for Phenolyzer and were scored based on previous biological knowledge regarding the specific cancer type. The cancer-specific Phenolyzer gene scores were used as a feature in the driver gene classification model.

### Membership to cancer-related pathways

The final features for the driver gene classification model were whether or not the gene is in the given selected cancer-related KEGG [25] pathways. The cancer-related KEGG pathways were determined as pathways related to the “Pathways in cancer” pathway: PPAR signaling pathway, MAPK signaling pathway, Calcium signaling pathway, cAMP signaling pathway, Cytokine-cytokine receptor interaction, HIF-1 signaling pathway, Cell cycle, p53 signaling pathway, mTOR signaling pathway, PI3K-Akt signaling pathway, Apoptosis, Wnt signaling pathway, Notch signaling pathway, Hedgehog signaling pathway, TGF-beta signaling pathway, VEGF signaling pathway, Focal adhesion, ECM-receptor interaction, Adherens junction, JAK-STAT signaling pathway, Estrogen signaling pathway.

For training the MTL model, the R package RMTL [50] was used. We trained an MTL model with sparse structure (lasso regularization). The framework for the algorithm used by RMTL is provided in Equation 1:

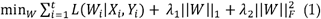

where *L*() is the logistic loss function. There are t tasks. *W* is the coefficient matrix, *W*_*i*_ is the *i*^th^ column of W and refers to the coefficient vector of task *i. X* is the predictor matrices and Y is the response vectors of the *t* tasks. ‖. ‖_1_ is the L_1_ norm and ‖. ‖_*F*_ is the Frobenius norm. *λ*_1_ aims to control the effect of cross-task regularization and *λ*_2_ stabilizes the numerical results and is used to improve generalization performance. Features for genes within the training data for all cancer types were further randomly split into training genes (75% of genes) and validation genes (25% of genes) datasets. The optimal *λ*_2_ value was obtained through the assessment of performance on the validation dataset (determined to be 10^−4^). After determining the optimal *λ*_1_ value using 10-fold cross-validation (determined to be 10^−4^), the final classification model was built.

For predicting the labels (i.e., “driver”, “non-driver”) using the probabilities of being a driver gene, cancer-type specific thresholds were determined as probability values maximizing accuracy on the validation datasets.

The performance of each sub-model was assessed by calculating AUC for each test dataset. Additionally, personalized analysis performances of the sub-models were assessed by calculating AUC per each individual in each test dataset.

### 2.3. Comparison with other batch analysis approaches

The performance of driveR was compared with the performances of the batch analysis approaches MutSigCV, DriverNet, OncodriveFML, and MutPanning by comparing AUC values.

MutSigCV (version 1.3.5) analyses were performed using default settings on the GenePattern platform [51]. DriverNet analyses were performed using DriverNet version 1.28.0 with the BioGRID [52, 53] Homo sapiens PIN (version 4.0.189) using the default settings. OncodriveFML analyses were performed using OncodriveFML version 2.3.0 with the default settings. MutPanning (version 2.0) analyses were performed using default settings on the GenePattern platform.

### 2.4. Comparison with other personalized analysis approaches

The personalized analysis performance of driveR was also compared with the performances of the personalized analysis approaches with DawnRank and PRODIGY by comparing AUC values.

As both of these tools required normal tissue expression data, only 16 datasets, for which more than 1 normal tissue expression data were available, were used, namely: BLCA-US, BRCA-US, CESC-US, COAD-US, HNSC-US, KIRC-US, KIRP-US, LIHC-US, LUAD-US, LUSC-US, PAAD-US, PRAD-US, READ-US, STAD-US, THCA-US, and UCEC-US.

DawnRank (version 1.2) analyses were performed with the BioGRID Homo sapiens PIN (version 4.0.189) using the default settings. PRODIGY (version 1.0) analyses were performed with the STRING [54] Homo sapiens PIN (version 11.0) and with KEGG curated pathways using the default settings. PRODIGY gene scores per pathway were aggregated into a single gene score for comparison purposes.

## 3. Results

### The variant impact metapredictor outperforms individual predictors

As described above, we trained a metapredictor model to estimate the pathogenicity of coding variants. Out of 6 different model types, the random forest metapredictor performed the best (Figure S2). As expected, this model also performed better than the individual impact prediction tools that were used as features to train the metapredictor, achieving an AUC of 0.911 (Figure 2).

**Figure 2.**
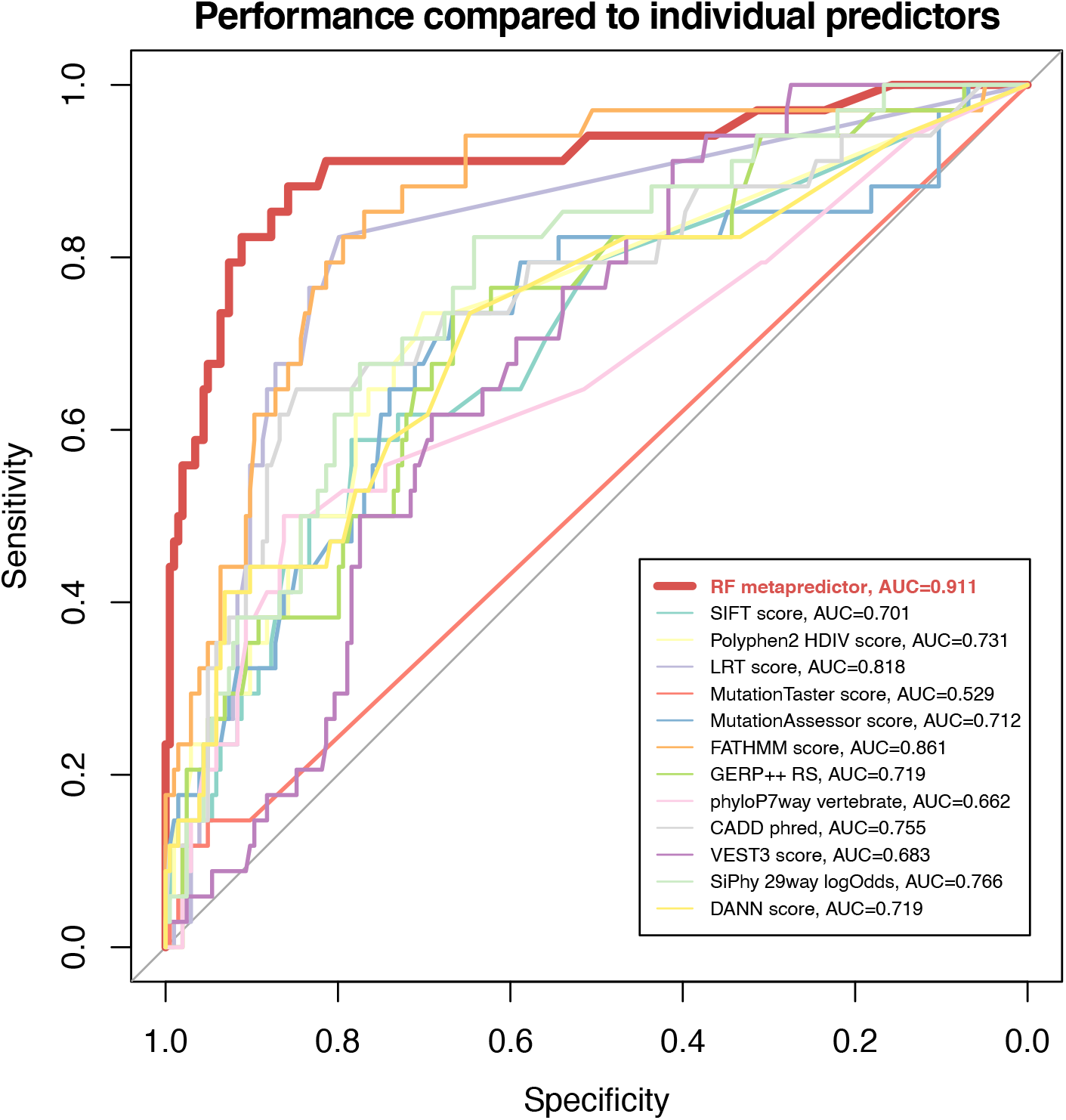
Metapredictor of coding variant impact outperforms individual predictors. ROC curves displaying the performances of the random forest (RF) metapredictor and individual impact prediction tools.

This metapredictor can be used to prioritize coding variants according to their pathogenicity and was used for generating one of the features of the MTL classification model.

### The cancer-type-specific driver gene prioritization approach performs well both for batch analysis and personalized analysis

Using genomic data and prior biological information, we next trained an MTL classification model for obtaining cancer-type-specific driver gene predictions. The batch analysis performance of the MTL classification model was assessed on the 28 test datasets (Figure 3). The median AUC on the test datasets was 0.684 (range=0.651-0.861, Figure 3A). As expected, the percentages of “True Driver Genes” (as curated by CGC) and “Actionable Genes” (as curated by TARGET [55], containing a total of 135 actionable genes) among all predicted driver genes across all datasets increased with increasing threshold values (Figure 3B, range of median percentages of “True Driver Genes” = 63.84-94.59%, range of median percentages of “Actionable Genes” = 55.29-85.71%). Using the cancer-type-specific thresholds (that maximized accuracy on the validation datasets), a median of 71.29% predicted driver genes were found to be “True Driver Genes” (Figure 3B left-hand panel), and 59.46% predicted driver genes were found to be “Actionable Genes” (Figure 3B right-hand panel).

**Figure 3.**
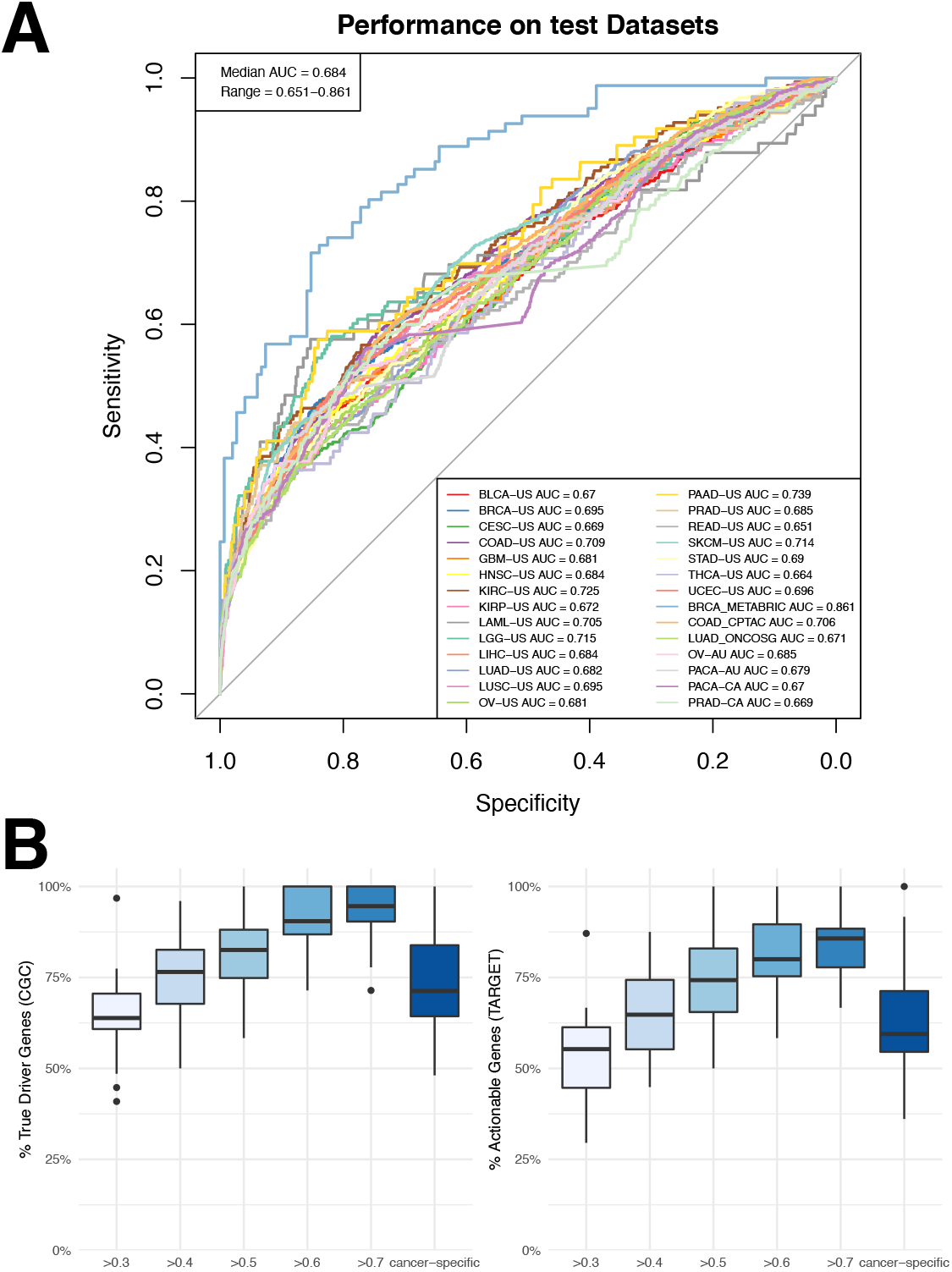
Batch analysis performance of driveR on 28 test datasets. (A) ROC curves of driveR prioritization results per each test dataset. AUC value per test dataset is indicated in the bottom-right legend. Median AUC and range are indicated on the top-left. (B) Boxplots of percentages of “True Driver Genes” (left) and “Actionable Genes” (right) among all predicted driver genes by using the specified thresholds across all datasets. “cancer-specific” indicates that the cancer-type-specific thresholds were used to predict driver genes.

Next, the personalized analysis performance of the driveR approach was assessed on the 5157 test patients (Figure 4). The median AUC value among all test patients was 0.773 (Figure 4A, top, range = 0-1). The median AUC values of patients per test dataset were similar (Figure 4A bottom, range of median AUC per dataset = 0.655-1). The median percentages of “True Driver Genes” and “Actionable Genes” among all predicted driver genes in each patient per all different thresholds were 100% (Figure 4B). Using the cancer-type specific thresholds, the median percentages were again 100%. This implies that each gene (if any) predicted by driveR to be a driver in an individual is most likely a true driver gene and an actionable gene.

**Figure 4.**
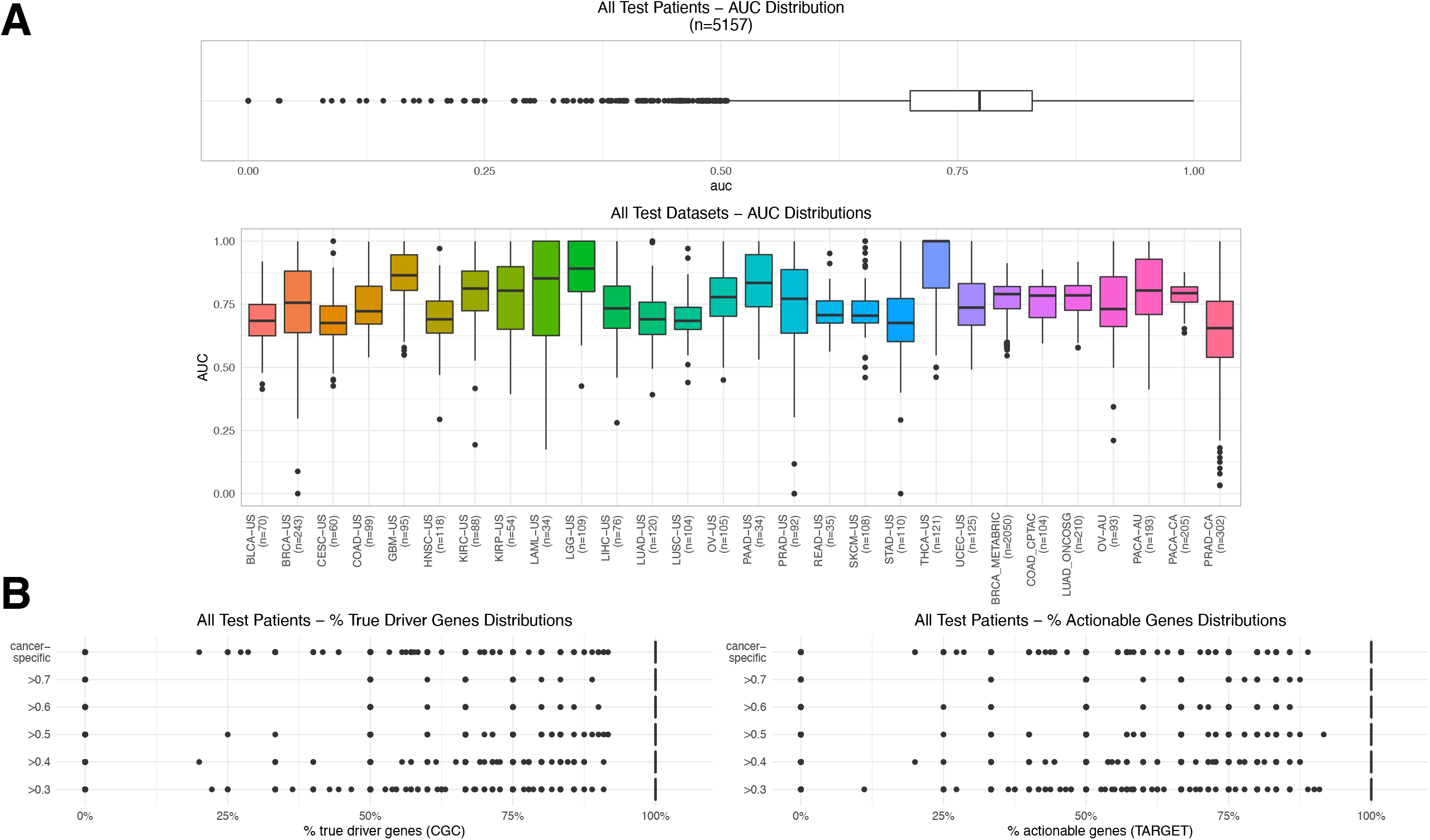
Personalized analysis performance of driveR on 5091 patients. (A) Boxplot displaying the distribution of AUC values in all patients (top). Boxplots displaying the distributions of AUC values per individual in each of the 28 test datasets. (B) Boxplots of percentages of “True Driver Genes” (left) and “Actionable Genes” (right) among all predicted driver genes by using the specified thresholds across all test patients. “cancer-specific” indicates that the cancer-type-specific thresholds were used to predict driver genes.

### The driver gene predictor approach outperforms existing approaches

The ROC curves of the performances of different batch analysis approaches per each dataset are presented in Figure S3. The median AUC of driveR across all test datasets (median AUC = 0.684) was significantly higher than all of MutSigCV (0.579, Wilcox rank sum test p < 0.001), DriverNet (0.614, p < 0.001), OncodriveFML (0.546, p < 0.001) and MutPanning (0.591, p < 0.001) (Figure 5).

**Figure 5.**
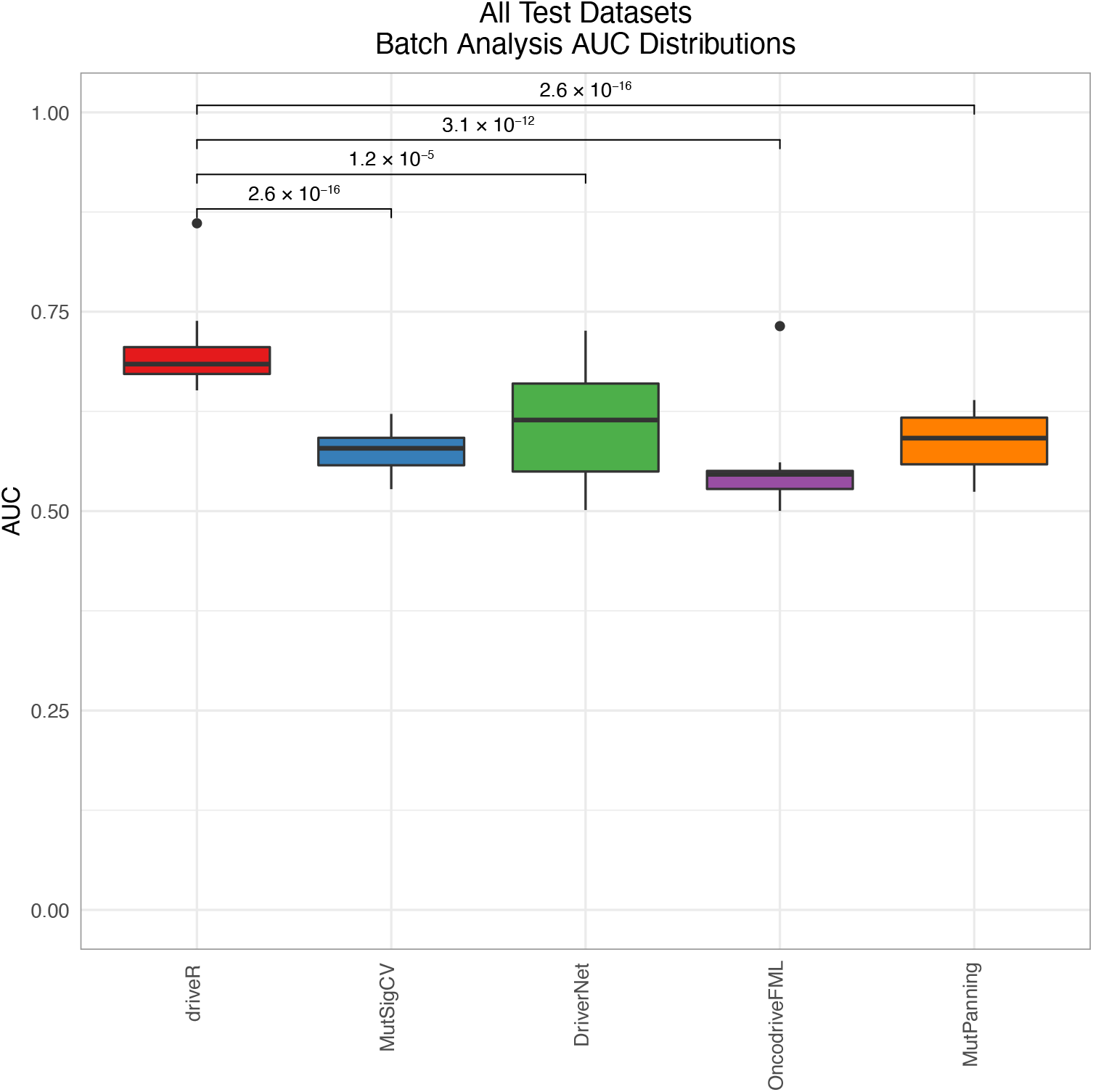
Comparison of performance of driveR with batch analysis approaches. Boxplots displaying the distributions of AUC values of each approach across the 28 test datasets. The brackets display p-values per each comparison.

Next, performances of driveR and other personalized analysis tools on test patients from 16 datasets, for which analyses with all approaches could be performed, were compared (Figure 6). It was observed that driveR had higher median AUC (0.728) compared to DawnRank (median AUC = 0.693, p < 0.001) and PRODIGY (median AUC = 0.679, p < 0.001) overall. When the performances were compared for all patients per dataset, driveR again displayed higher performance (Figure S4).

**Figure 6.**
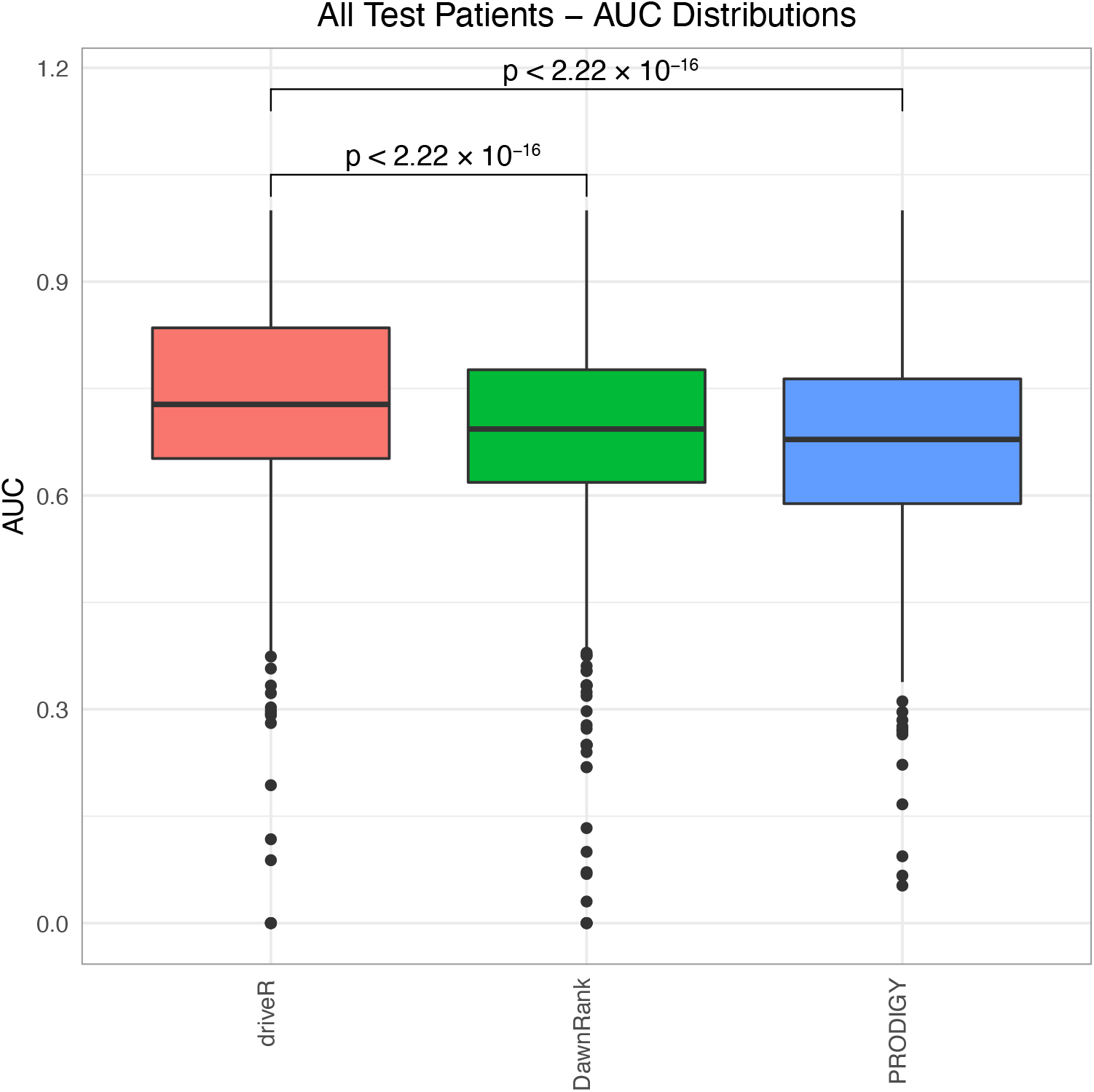
Comparison of performance of driveR with personalized analysis approaches. Boxplots displaying the distributions of AUC values of each approach across all test patients. The brackets display p-values per each comparison.

## 4. Discussion

Alterations in driver genes are the putative underlying causes of oncogenesis and tumor formation [3, 4]. Although numerous driver genes are experimentally validated [56-59], there is great potential clinical benefit in identifying driver genes in individual patients [5, 6].

In this study, we established a simple, model-based approach, driveR, that can accurately prioritize cancer driver genes, sorting through a vast amount of passengers observed in cancer genomes [1]. Using 26 features based on somatic genomics data, we trained a multi-task learning classification model to estimate driver gene probabilities in a cancer-type-specific context for 21 different cancer types. Compared to other approaches, driveR achieved better performance on different test datasets by accurately prioritizing cancer driver genes in analyses on both cohort-scale and personalized data. Below we discuss several unique aspects of driveR.

Driver genes are diverse among different cancer types [56-59]. Different cancers may possess different driver genes. The driveR approach attempts to define cancer-type-specific driver genes based on somatic genomic features, incorporating prior biological knowledge. The multi-task learning model at the core of our approach allows for cancer-type-specific identification of driver genes. The approach is available for use for 21 different cancer types. Analysis in a cancer-type-specific context allows driveR to more accurately identify driver genes that may be specific to a certain cancer type.

Most approaches for driver gene prioritization are designed for analyzing tumor cohorts. These approaches usually fail to identify low-frequency driver genes, not to mention patient-specific driver genes. Patient-specific driver genes may be rare or not match the organ-of-origin. In this study, we demonstrate that driveR can be utilized to rapidly and accurately analyze both patient-specific and cohort-scale genomic data to prioritize cancer driver genes. This makes driveR a suitable option for studying driver genes for individual patients.

In our approach, we incorporated prior biological knowledge into the MTL model. We used Phenolyzer, a database-mining tool, which integrates 15 different biological knowledge databases to score a gene’s prior association with cancer. Additionally, memberships of genes to cancer-related KEGG pathways were also taken into consideration. This integration of extensive biological knowledge improved the accuracy of the MTL model because the final model used for driver gene prioritization was based not only on the genomic features but also guided by expert knowledge from decades of research.

We devised driveR to be less demanding on the data type. Other driver prioritization tools usually require multiple omics data for the patient, including genomic mutation data, tumor expression data, and normal expression data. However, obtaining transcriptomics data is not always feasible due to cost and other practical issues. We devised driveR based only on somatic genomics data (somatic mutation and SCNA) because targeted sequencing or whole exome/genome sequencing is more widely utilized in both the clinical and research settings and technical analysis of genomic data is less complicated compared to transcriptomic data.

We demonstrated that the overall performance of driveR in identifying true drivers is adequate on both batch analyses and personalized analyses. Additionally, a high proportion of driveR predictions were also clinically-actionable genes. We also demonstrate that driveR outperforms other tools in both personalized and batch analyses. These demonstrate that driveR performs well in successfully prioritizing driver genes in a cancer-type-specific context.

## 5. Conclusions

In this study, we devised a novel approach, driveR, for prioritizing cancer-type-specific driver genes using somatic genomic data. As demonstrated, driveR can be utilized for both analyzing individual cancers and cancer cohorts to accurately prioritize patient-specific or cohort-scale driver genes. We also demonstrated that our approach outperforms existing driver gene prioritization methods. We hope that this approach can provide further insight into cancer driver gene discovery and help progress personalized cancer research.

## 6. List of abbreviations

PIN: Protein-protein interaction network
SCS: Single-sample controller strategy
DEG: differentially-expressed gene
KEGG: Kyoto Encyclopedia of Genes and Genomes
SVM: Support Vector Machine
ROC: Receiver Operating Characteristic
AUC: Area Under Curve
MTL: Multi-task learning
SCNA: Somatic Copy Number Alteration
TCGA: The Cancer Genome Atlas
ICGC: International Cancer Genome Consortium
CGC: Cancer Gene Census
MCR: Minimal Common Region
COSMIC: Catalogue of Somatic Mutations in Cancer
Phenolyzer: Phenotype Based Gene Analyzer

## 7. Declarations

### Ethics approval and consent to participate

Not applicable

### Consent for publication

Not applicable

### Availability of data and materials

The datasets used and/or analyzed during the current study are available from the corresponding author on reasonable request.

Project name: driveR

Project home page: https://github.com/egeulgen/driveR

Operating system(s): Platform independent

Programming language: R

License: MIT license

### Authors’ contributions

Conceptualization, EU and OUS; implementation, analyses, preparing the manuscript draft, EU; study supervision, OUS; review and editing, both authors. Both authors have read and agreed to the published version of the manuscript.

## Acknowledgements

Not applicable

## 8. Figure Legends

**Figure S1.**
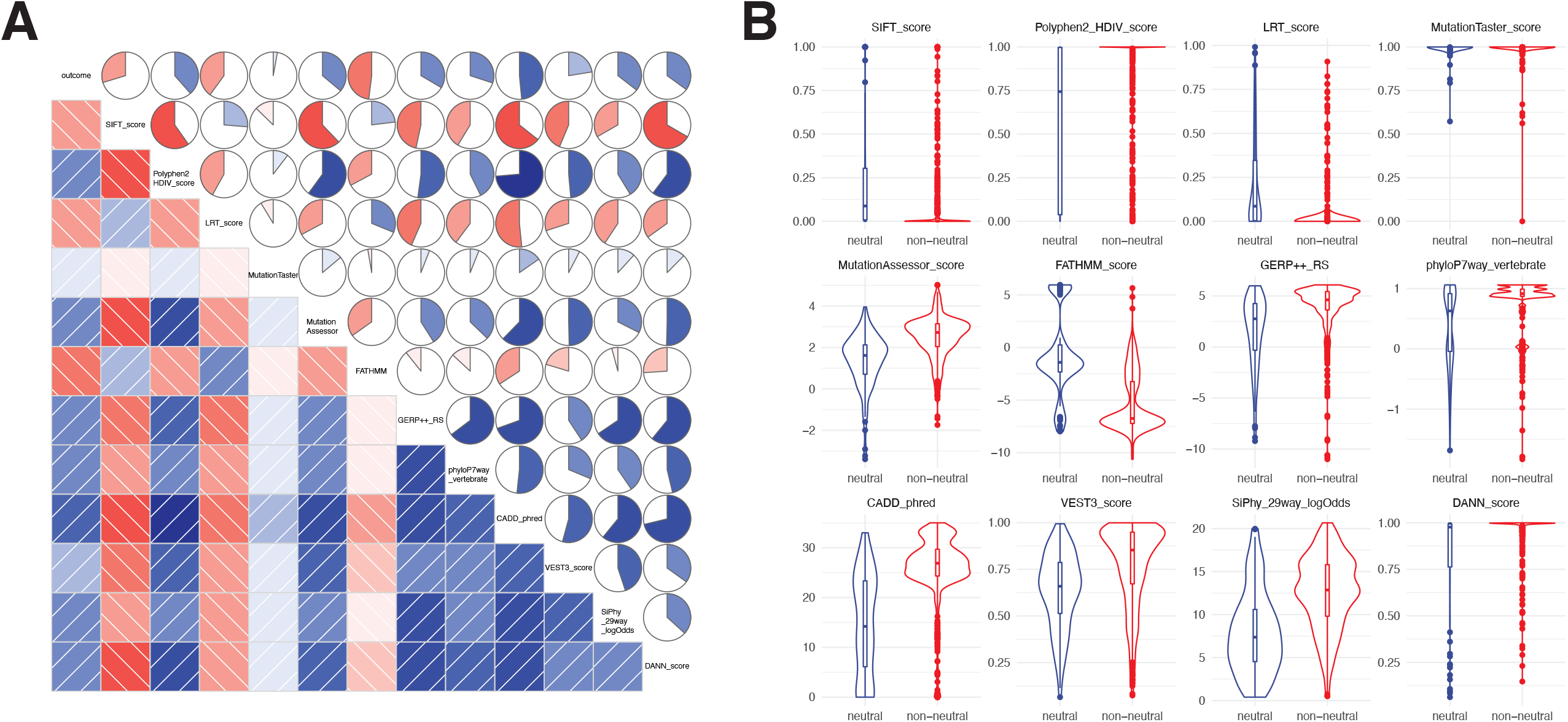
The overall analysis of individual variant impact predictors. (A) Correlogram displaying correlations between the outcome and individual variant impact predictors. (B) Violin plots displaying the distributions of scores of individual variant impact predictors in driver and non-driver variants.

**Figure S2.**
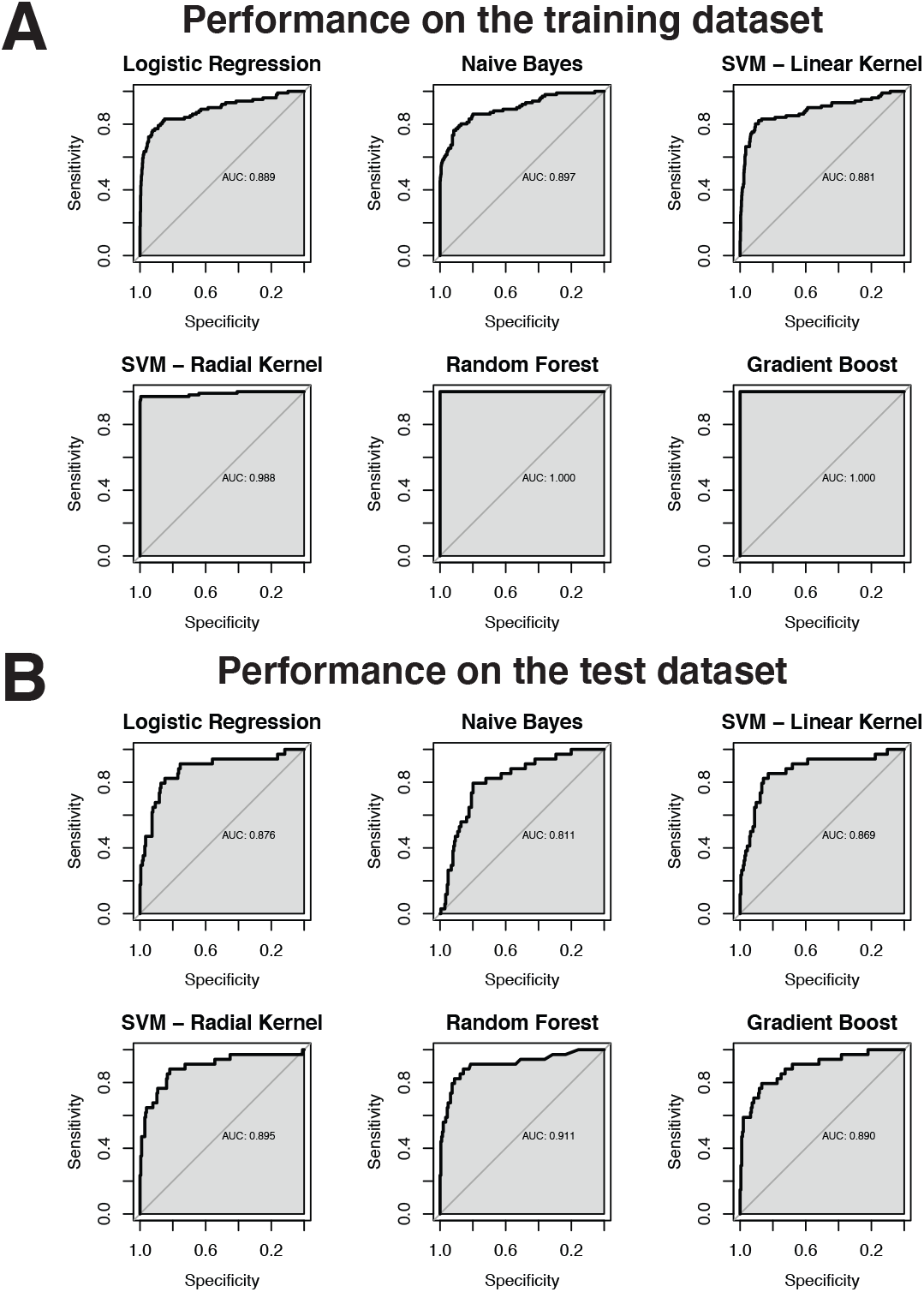
Performance of different coding variant impact metapredictor models in the training (A) and test (B) datasets

**Figure S3.**
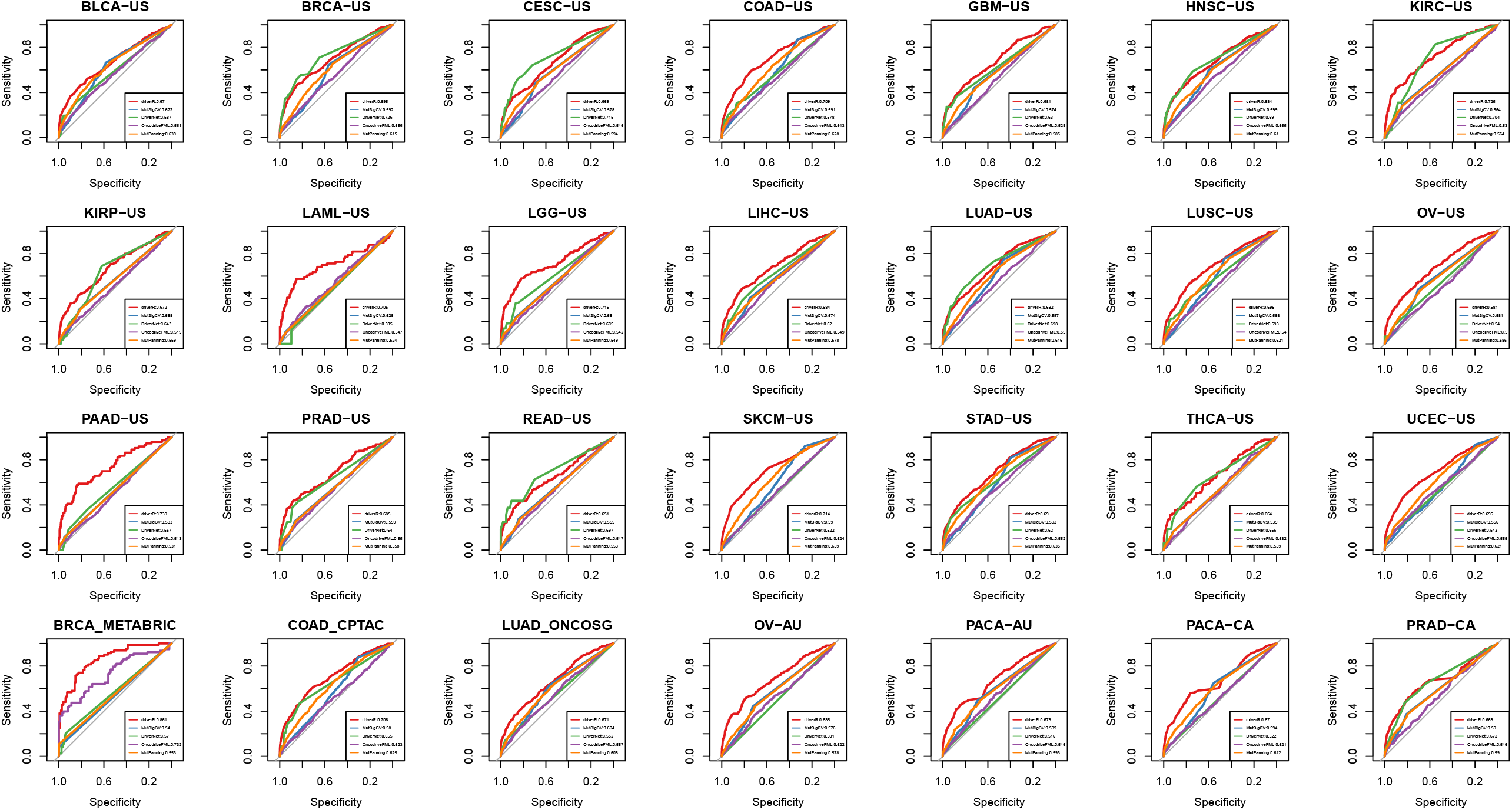
Comparison of performance of driveR with batch analysis approaches per test dataset. ROC curves for assessing the performance of each approach per each test dataset. The bottom-right legends display AUC per each approach.

**Figure S4.**
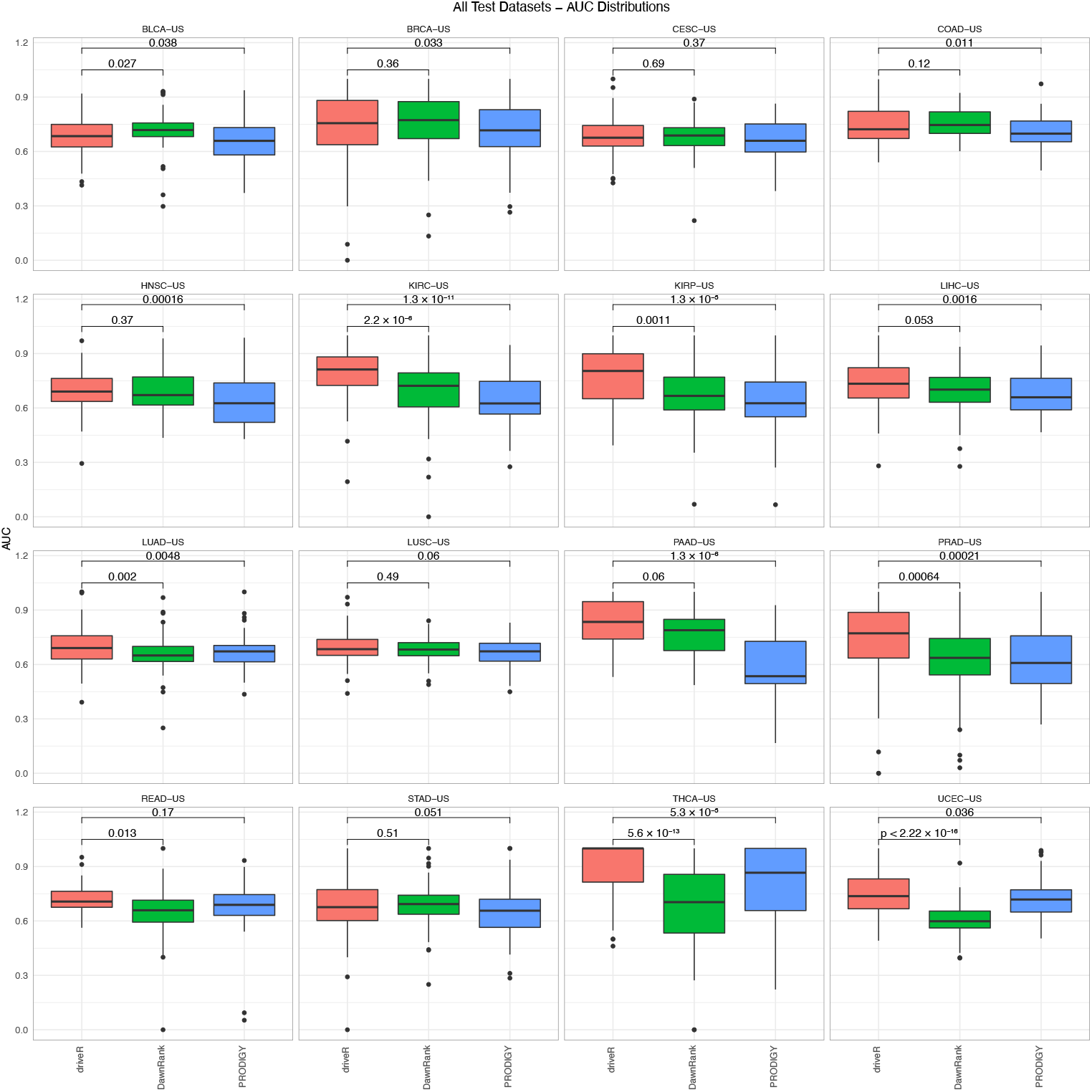
Comparison of performance of driveR with personalized analysis approaches per test dataset. Boxplots displaying the distributions of AUC values of each approach across all patients per test dataset. The bracket display p values per each comparison.

