## Supplementary Table 1 for "driveR: A Novel Method for Prioritizing Cancer Driver Genes Using Somatic Genomics Data"

**Supplementary Table 1.** Datasets used for training and testing the multi-task learning classification model. “Type” indicates whether the dataset was used for training or testing. “Dataset ID” indicates the name of the dataset. “Cancer type” indicates the cancer type of the dataset. “Source” indicates the source from which the dataset was obtained. “Original Source” indicates the original source of the data.

| Type | Dataset ID | Cancer Type | Source | Original Source |
| --- | --- | --- | --- | --- |
| Test | BLCA-US | Bladder Urothelial Cancer | ICGC Data Portal Release 28 | TCGA, US |
| Test | BRCA_METABRIC | Breast Cancer | cBioPortal - Datasets | METABRIC |
| Test | BRCA-US | Breast Cancer | ICGC Data Portal Release 28 | TCGA, US |
| Test | CESC-US | Cervical Squamous Cell Carcinoma | ICGC Data Portal Release 28 | TCGA, US |
| Test | COAD_CPTAC | Colon Adenocarcinoma | cBioPortal - Datasets | CPTAC-2 Prospective |
| Test | COAD-US | Colon Adenocarcinoma | ICGC Data Portal Release 28 | TCGA, US |
| Test | GBM-US | Brain Glioblastoma Multiforme | ICGC Data Portal Release 28 | TCGA, US |
| Test | HNSC-US | Head and Neck Squamous Cell Carcinoma | ICGC Data Portal Release 28 | TCGA, US |
| Test | KIRC-US | Kidney Renal Clear Cell Carcinoma | ICGC Data Portal Release 28 | TCGA, US |
| Test | KIRP-US | Kidney Renal Papillary Cell Carcinoma | ICGC Data Portal Release 28 | TCGA, US |
| Test | LAML-US | Acute Myeloid Leukemia | ICGC Data Portal Release 28 | TCGA, US |
| Test | LGG-US | Brain Lower Grade Glioma | ICGC Data Portal Release 28 | TCGA, US |
| Test | LIHC-US | Liver Hepatocellular carcinoma | ICGC Data Portal Release 28 | TCGA, US |
| Test | LUAD_ONCOSG | Lung Adenocarcinoma | cBioPortal - Datasets | OncoSG |
| Test | LUAD-US | Lung Adenocarcinoma | ICGC Data Portal Release 28 | TCGA, US |
| Test | LUSC-US | Lung Squamous Cell Carcinoma | ICGC Data Portal Release 28 | TCGA, US |
| Test | OV-AU | Ovarian Cancer | ICGC Data Portal Release 28 | AU |
| Test | OV-US | Ovarian Serous Cystadenocarcinoma | ICGC Data Portal Release 28 | TCGA, US |
| Test | PAAD-US | Pancreatic Cancer | ICGC Data Portal Release 28 | TCGA, US |
| Test | PACA-AU | Pancreatic Cancer | ICGC Data Portal Release 28 | AU |
| Test | PACA-CA | Pancreatic Cancer | ICGC Data Portal Release 28 | CA |
| Test | PRAD-CA | Prostate Adenocarcinoma | ICGC Data Portal Release 28 | CA |
| Test | PRAD-US | Prostate Adenocarcinoma | ICGC Data Portal Release 28 | TCGA, US |
| Test | READ-US | Rectum Adenocarcinoma | ICGC Data Portal Release 28 | TCGA, US |
| Test | SKCM-US | Skin Cutaneous melanoma | ICGC Data Portal Release 28 | TCGA, US |
| Test | STAD-US | Gastric Adenocarcinoma | ICGC Data Portal Release 28 | TCGA, US |
| Test | THCA-US | Head and Neck Thyroid Carcinoma | ICGC Data Portal Release 28 | TCGA, US |
| Test | UCEC-US | Uterine Corpus Endometrial Carcinoma | ICGC Data Portal Release 28 | TCGA, US |
| Training | BLCA-US | Bladder Urothelial Cancer | ICGC Data Portal Release 28 | TCGA, US |
| Training | BRCA-US | Breast Cancer | ICGC Data Portal Release 28 | TCGA, US |
| Training | CESC-US | Cervical Squamous Cell Carcinoma | ICGC Data Portal Release 28 | TCGA, US |
| Training | COAD-US | Colon Adenocarcinoma | ICGC Data Portal Release 28 | TCGA, US |
| Training | GBM-US | Brain Glioblastoma Multiforme | ICGC Data Portal Release 28 | TCGA, US |
| Training | HNSC-US | Head and Neck Squamous Cell Carcinoma | ICGC Data Portal Release 28 | TCGA, US |
| Training | KIRC-US | Kidney Renal Clear Cell Carcinoma | ICGC Data Portal Release 28 | TCGA, US |
| Training | KIRP-US | Kidney Renal Papillary Cell Carcinoma | ICGC Data Portal Release 28 | TCGA, US |
| Training | LAML-US | Acute Myeloid Leukemia | ICGC Data Portal Release 28 | TCGA, US |
| Training | LGG-US | Brain Lower Grade Glioma | ICGC Data Portal Release 28 | TCGA, US |
| Training | LIHC-US | Liver Hepatocellular carcinoma | ICGC Data Portal Release 28 | TCGA, US |
| Training | LUAD-US | Lung Adenocarcinoma | ICGC Data Portal Release 28 | TCGA, US |
| Training | LUSC-US | Lung Squamous Cell Carcinoma | ICGC Data Portal Release 28 | TCGA, US |
| Training | OV-US | Ovarian Serous Cystadenocarcinoma | ICGC Data Portal Release 28 | TCGA, US |
| Training | PAAD-US | Pancreatic Cancer | ICGC Data Portal Release 28 | TCGA, US |
| Training | PRAD-US | Prostate Adenocarcinoma | ICGC Data Portal Release 28 | TCGA, US |
| Training | READ-US | Rectum Adenocarcinoma | ICGC Data Portal Release 28 | TCGA, US |
| Training | SKCM-US | Skin Cutaneous melanoma | ICGC Data Portal Release 28 | TCGA, US |
| Training | STAD-US | Gastric Adenocarcinoma | ICGC Data Portal Release 28 | TCGA, US |
| Training | THCA-US | Head and Neck Thyroid Carcinoma | ICGC Data Portal Release 28 | TCGA, US |
| Training | UCEC-US | Uterine Corpus Endometrial Carcinoma | ICGC Data Portal Release 28 | TCGA, US |
